# Complex-specific subunits of endosomal tethering factors determine the balance between HOPS/CORVET complexes

**DOI:** 10.1101/2023.08.31.555722

**Authors:** Ármin Sőth, Márton Molnár, Péter Lőrincz, Zsófia Simon-Vecsei, Gábor Juhász

## Abstract

The endolysosomal tethering complexes HOPS and CORVET play pivotal roles in the homo- and heterotypic fusion of early and late endosomes, respectively, and HOPS also mediates the fusion of lysosomes with incoming vesicles including late endosomes and autophagosomes. These heterohexameric complexes share their four core subunits that assemble with additional two, complex-specific subunits. These features and the similar structure of the complexes allow the formation of hybrid complexes, and the complex specific subunits may compete for binding to the core. In our study, we decided to gain insight into how human HOPS and CORVET complexes form. We found that the overexpression of CORVET-specific Vps8 or Tgfbrap1 decreased the amount of core proteins Vps11 and Vps18 that are assembled with HOPS-specific subunits Vps41 or Vps39, which suggests reduced amount of intracellular HOPS. In line with this, we observed that the level of lipidated LC3 protein was elevated in these cells and the autophagic cargo p62 showed accumulation in cells overexpressing Vps8, suggesting failure in autophagosome-lysosome fusion in line with loss of HOPS function. In contrast, overexpression of HOPS-specific Vps39 or Vps41 did not affect the levels of these autophagic markers. Finally, we found that a hybrid complex containing Vps39 and Vps8 could be formed in HEK293 cells.

## Introduction

Living organisms require coordinated operation of biosynthetic and degradative pathways for maintaining cellular homeostasis. To achieve this, eukaryotic cells sustain various membrane-bound organelles, and proteins and lipids are constantly exchanged between them via vesicle transport. This process depends on multiple fusion and fission events. Vesicular fusion is executed by a conserved machinery consisting of Rab GTPases, their interacting effectors and SNARE proteins that are found on both membranes (Bonifacino and Glick 2004). The maturation of endosomes is regulated by Rab proteins, and their GTP-bound, active form can bind multiple effectors, such as tethering factors. These can bring the donor and acceptor membranes into contact and thus facilitate the homo- and heterotypic fusion of vesicles (Bröcker et al. 2012). The fusion is completed by SNAREs that mediate the mixing of the lipid bilayers (Bonifacino and Glick 2004).

Endolysosomal tethering complexes, namely CORVET (Class C core endosome vacuole tethering) and HOPS (Homotypic vacuole fusion and protein sorting) were first identified in yeast (Seals et al. 2000); (Wurmser et al. 2000; Peplowska et al. 2007). The homotypic fusion of Vps21/Rab5-positive endosomes is mediated by CORVET (Balderhaar et al. 2013), whilst HOPS promotes the fusion of late endosomes (Ypt7/Rab7-positive) and autophagosome-lysosome fusion as well (Angers and Merz, 2009; Balderhaar and Ungermann 2013; Takáts et al. 2014). In yeast, the complexes consist of four shared, core subunits (Vps16, Vps33, Vps18, Vps11), and two complex-specific subunits: Vps8 and Vps3 for CORVET (Peplowska et al. 2007) and Vps41 and Vps39 for HOPS (Seals et al. 2000); (Wurmser et al. 2000). The specific subunits are responsible for targeting the complexes to specific membranes by binding to membrane-bound (thus GTP loaded) Rab small GTPases (Rab2 and Rab7 in the case of HOPS and Rab5 for CORVET), and hence they bring the fusion-primed vesicles closer to each other.

In Drosophila, a tetrameric miniCORVET complex was identified, which lacks Vps11 and Vps3 (the latter one has no homologs in higher eukaryotes) (Lőrincz et al. 2016). In mammals, CORVET harbours Tgfbrap1 instead of Vps3 (Lachmann et al. 2014; Perini et al. 2014), whilst HOPS is conserved in metazoans.

The existence of shared subunits raises the possibility of interconversion between HOPS and CORVET, and hence the formation of hybrid complexes. Biochemical experiments in yeast identified complexes containing the class C Vps proteins (Vps16, Vps33, Vps18 and Vps11) and one CORVET- and one HOPS-specific subunits: “i-HOPS”: Vps8-Vps39 and “i-CORVET”: Vps41-Vps3 (Peplowska et al. 2007; Ostrowicz et al. 2010). However, in Drosophila, the only possible hybrid complex (Vps8-Vps39) could not be found based on mass spectrometry data showing interacting proteins of CORVET-specific Vps8 (Lőrincz et al. 2016). To date, there are no data about the existence of mammalian hybrid complexes.

Furthermore, the shared class C proteins and overlapping binding sites (Ostrowicz et al. 2010; Van Der Kant et al. 2015) allows potential competition between the specific subunits, which can affect intracellular levels of the complexes. In yeast, Vps3 overexpression decreases the level of assembled HOPS and leads to vacuole fragmentation, which resembles to the Vps39 (HOPS) loss of function phenotype. However, overproduction of Vps8 did not have such effect (Peplowska et al. 2007). In contrast, overexpression of Drosophila Vps8, the only miniCORVET-specific subunit, inhibits HOPS-dependent trafficking routes by outcompeting Vps41 from HOPS, while increased amount of HOPS-specific Vps41 does not disturb CORVET functions (Lőrincz et al. 2019). Based on an electron microscopy study revealing the structure of the yeast HOPS complex (Bröcker et al. 2012) and detailed characterization of the architecture of mammalian complexes (Van Der Kant et al. 2015), it is likely that similar competition can occur between the specific subunits of the human complexes as well. However, this question has not been investigated yet in mammalian cells.

In our study, we show that overexpression of human CORVET-specific Vps8 or Tgfbrap1 decreased the intracellular level of HOPS, while the gain of their HOPS-specific counterparts, Vps41 and Vps39, respectively, did not affect CORVET. Additionally, high level of Vps8 increased lipidated LC3 and p62 levels, in line with decreased HOPS function. We also show the possible formation of human Vps39-Vps8 hybrid complex.

## Results

### Overexpression of CORVET-specific subunits decreased the amount of HOPS

Human HOPS and CORVET are heterohexamers, similarly to the yeast complexes, where overexpression of Vps3 can interfere with Vps39 assembling into HOPS (Peplowska et al. 2007). This competition model is based on formation of more Vps41-Vps3 intermedier complexes, and less HOPS (Vps41-Vps39) because of elevated Vps3 level. Vps8 overexpression decreases the amount of Vps41-Vps3 hybrid complex, however, the amount of HOPS seems to be unaffected (Peplowska et al. 2007).

In Drosophila, a smaller CORVET complex exists with only one complex-specific subunit: Vps8 (Lőrincz et al. 2016). As a result of its overexpression, the amount of HOPS decreased based on immunoprecipitation (IP) data and functional tests (Lőrincz et al. 2019). In flies, the effect of the other CORVET-specific subunit could not be investigated, but both CORVET-specific subunits can be examined in human cell cultures.

First we investigated the effect of CORVET-specific subunits on the intracellular level of HOPS complex. Tgfbrap1 and Vps8 were overexpressed one by one in HEK293 cells stably transfected with Vps39-FLAG or Vps41-FLAG, respectively, and after anti-FLAG IP, the amount of core subunits Vps11 and Vps18 were analyzed. Vps41 could precipitate 61% less Vps11 and 47% less Vps18 upon Vps8 overproduction compared to control cells (Fig.1A). Similarly, Vps39 could bind lower amount of Vps11 (decreased by 24%) and Vps18 (decreased by 21%) when Tgfbrap1 was overexpressed (Fig.1B). These data suggest that in excess amount, CORVET-specific subunits successfullycompete with their HOPS-counterparts, and hence less core components can be assembled with HOPS-specific subunits.

**Figure 1.**
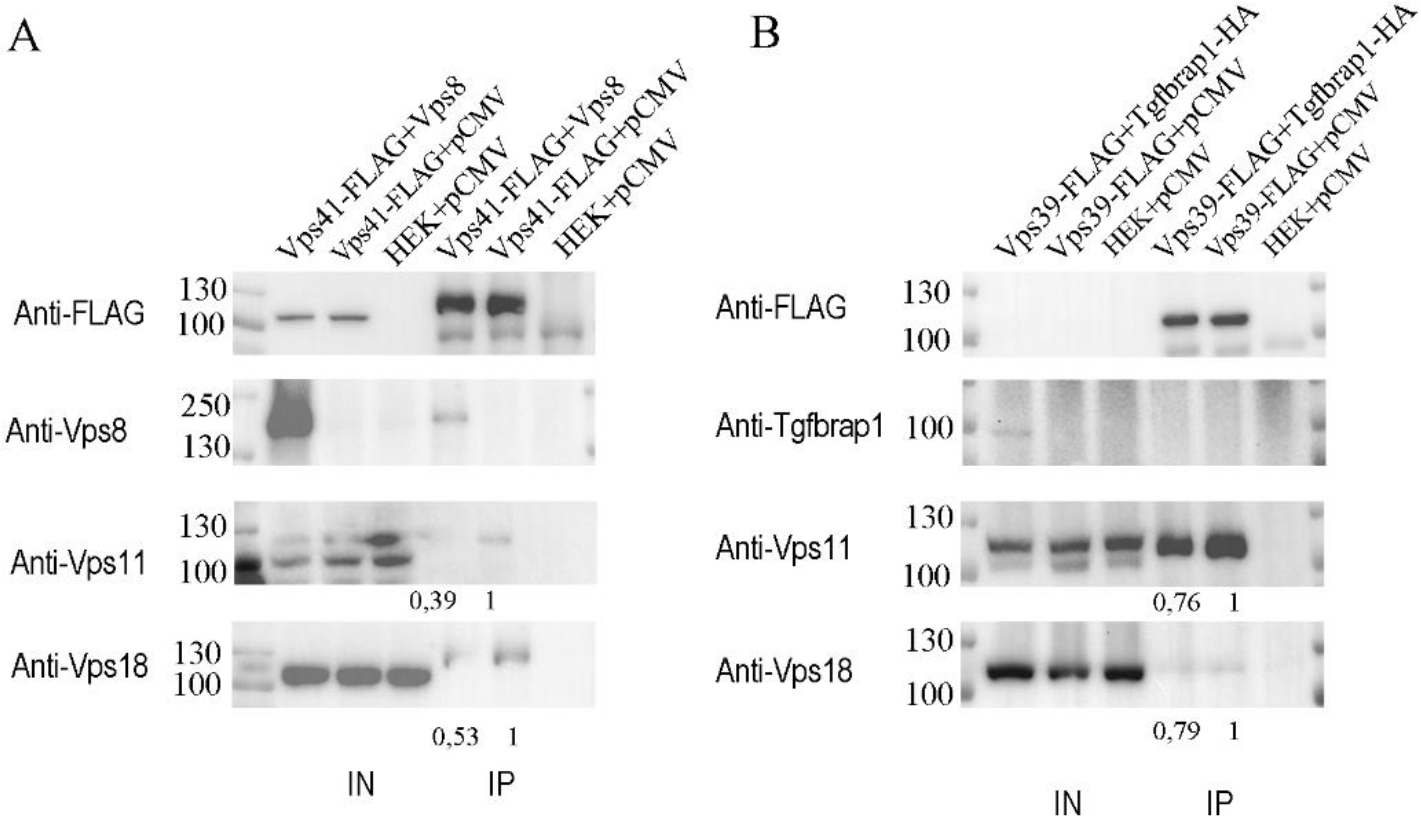
The overproduction of Vps8 or Tgfbrap1 decreases the amount of HOPS-bound Vps11 and Vps18. HEK293 cells were stably transfected with Vps41-FLAG (A) or Vps39-FLAG (B) and transiently transfected with Vps8 (A) or Tgfbrap1 (B), respectively. Total cell lysates were applied on anti-FLAG resin and bound proteins were detected by western blot. Overexpression of CORVET-specific subunits decreases the amounts of Vps41- (A) or Vps39-bound (B) core subunits (Vps11 and Vps18) compared to control cells (HEK + pCMV). Relative amounts of Vps41-(A) or Vps39-bound (B) core subunits were calculated by densitometry using Image J and are shown below each panel. Representative figures of at least three experiments.

### Overexpression of HOPS-specific subunits did not affect the amount of CORVET

Next, we determined the effect of HOPS-specific subunit overexpression on CORVET. We followed the same logic as above and overexpressed Vps39 and Vps41 in cells stably transfected with Tgfbrap1-FLAG or Vps8-FLAG, respectively. After anti-FLAG IP, we found no change in the amount of bound core subunits, Vps11 and Vps18, despite the Vps39 or Vps41 overexpression (Fig. 2A and B). These indicated that elevated amount of HOPS-specific subunits could not outcompete CORVET-specific Tgfbrap1 and Vps8, and hence the level of assembled CORVET is not affected.

**Figure 2.**
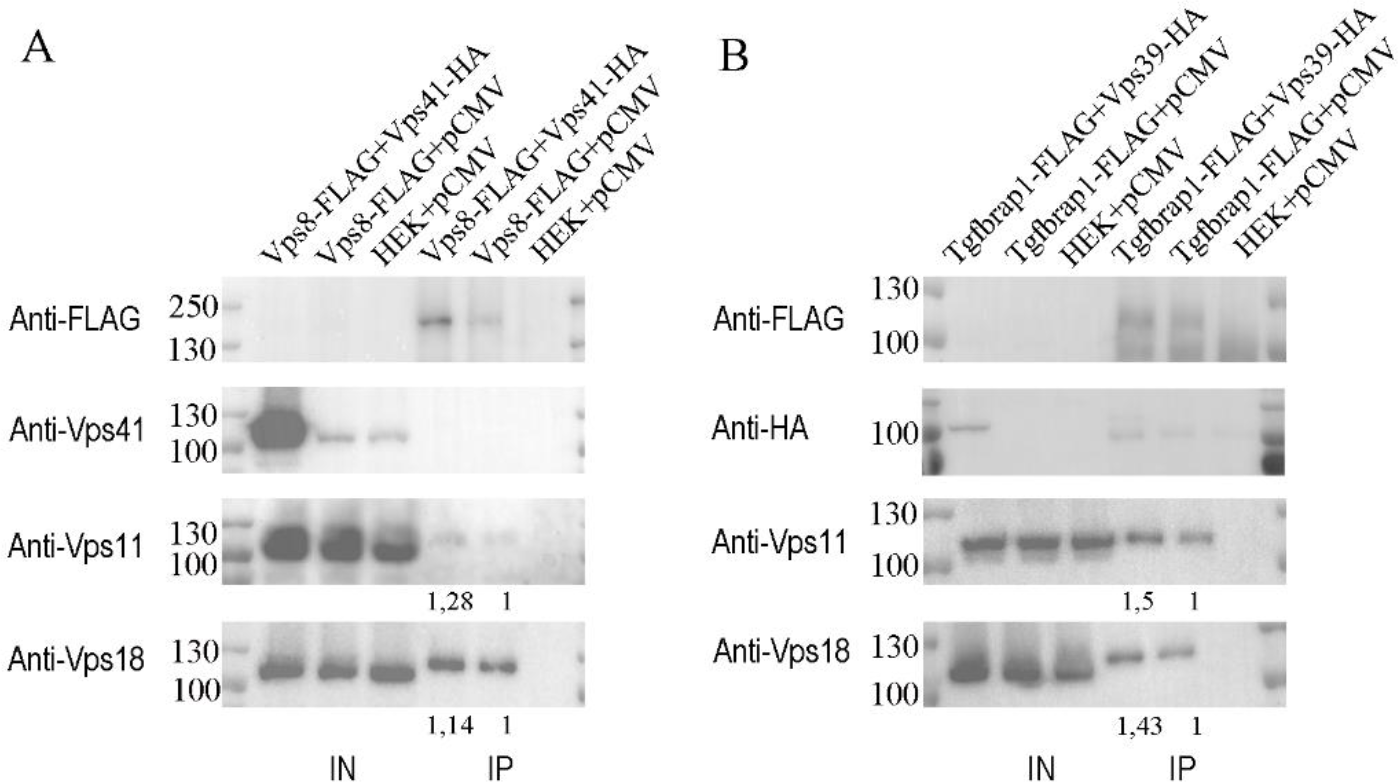
The overproduction of Vps39 or Vps41 does not affect the amount of CORVET-bound Vps11 and Vps18. HEK293 cells were stably transfected with Vps8-FLAG (A) or Tgfbrap1-FLAG (B) and transiently transfected with Vps41-HA (A) or Vps39-HA (B), respectively. Total cell lysates were applied on anti-FLAG resin and bound proteins were detected by western blot. Overexpression of HOPS-specific subunits did not change the amounts of Vps8- (A) or Tgfbrap1-bound (B) core subunits (Vps11 and Vps18) compared to control cells (HEK + pCMV). Relative amounts of Vps8-(A) or Tgfbrap1-bound (B) core subunits were calculated by densitometry using Image J and are shown below each panel. Representative figures of at least three experiments.

### Overexpression of CORVET-specific subunits increased the amount of lipidated LC3 and p62

During macroautophagy, autophagosomes transfer their cargo to the lysosomal compartment by fusion with lysosomes. HOPS mediates autophagosome-lysosome fusion (Angers and Merz 2009; Balderhaar and Ungermann 2013; Takáts et al. 2014), besides the homo- and heterotypic fusions of late endosomes. To assess the effect of CORVET-specific subunit overexpression on HOPS function, we examined later steps of the autophagic processes using different methods. Accumulation of the lipid-bound form of microtubule-associated protein 1A/1B light chain 3B (LC3) or the autophagic cargo p62 can indicate the failure of autophagosome-lysosome fusion as a readout of HOPS function (Jiang et al. 2014).

To examine the effect of the overexpressed specific subunits on HOPS function, we used our HEK293 cell lines stably transfected with FLAG-tagged versions of the different subunits. Intracellular levels of LC3 and p62 were analyzed by western blot (WB). Cells overexpressing CORVET-specific Tgfbrap1 or Vps8 showed elevated lipidated LC3 (LC3-II) level, similarly to cells lacking the core subunit Vps18 (Vps18KO; both HOPS and CORVET functions are impaired). HOPS-specific subunits Vps39 and Vps41 did not have the same effect (Fig.3A and B). Additionally, the level of autophagic cargo p62 increased in cells overproducing Vps8 and in Vps18KO cells, however, the other subunits did not elevate the amount of p62 (Fig.3A and B).

**Figure 3.**
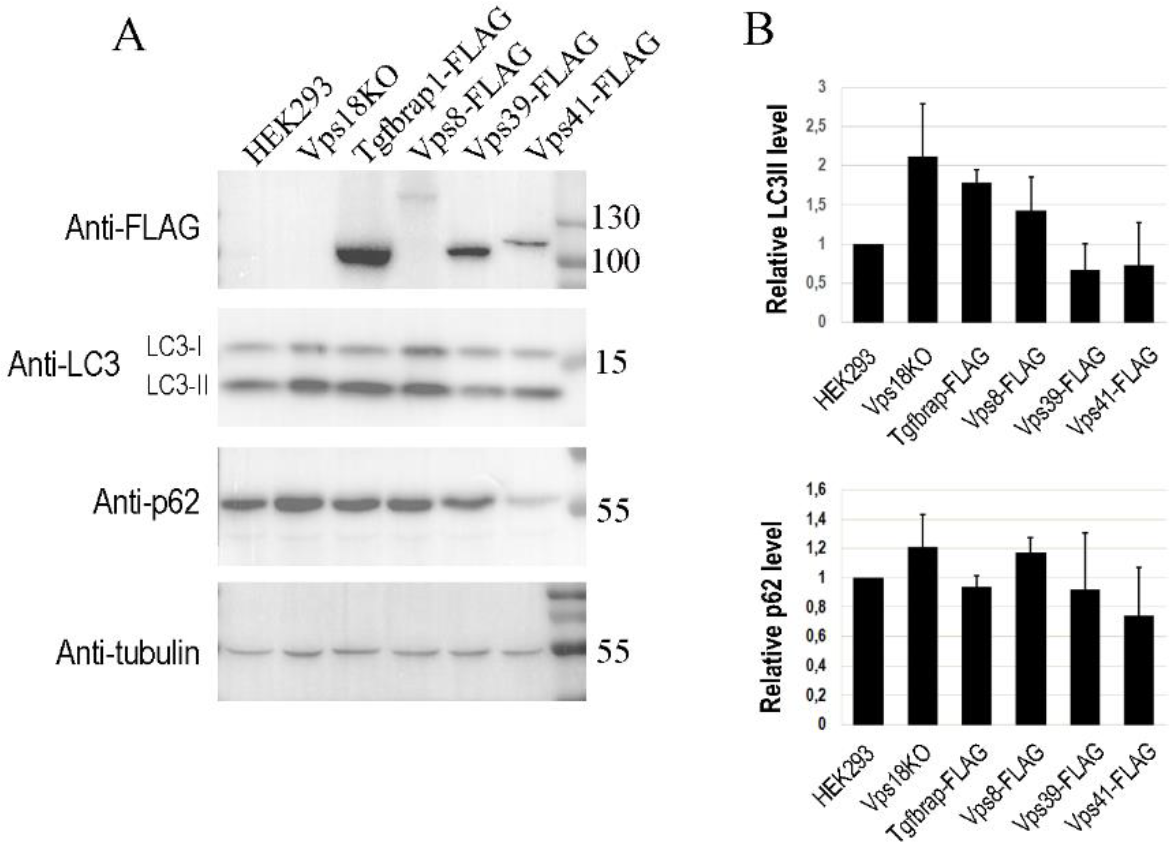
Overexpression of Vps8 interferes with autophagy. Total cell lysates of Vps18 KO and HEK293 overexpressing different FLAG-tagged HOPS- or CORVET subunits were investigated by western blot. The levels of the overexpressed proteins (anti-FLAG), LC3, p62 and tubulin (loading control) were detected (A). Vps8 overexpression elevated the level of both lipidated LC3 (LC3II) and the autophagic cargo p62, while the surplus of Tgfbrap1 caused LC3II accumulation. The quantification of LC3 and p62 blots were done by densitometry using Image J, based on three experiments (B).

Taken together we detected elevated LC3-II and p62 levels in cells with surplus of Vps8, which suggests altered HOPS function.

### Hybrid complex can be formed that contains HOPS-specific Vps39 and CORVET-specific Vps8

Yeast data indicates that hybrid complexes can be formed: Vam6/Vps39 - Vps8 complex could be purified through TAP-tagged Vam6/Vps39, however, its amount was much less than the “wild-type” complexes. A small amount of Vps41 - Vps3 complex was also identified in control cells, while its level was elevated in cells lacking Vps8 or Vam6/Vps39 (Peplowska et al. 2007; Ostrowicz et al. 2010). In Drosophila, Vps8 is the only CORVET-specific subunit, hence only the existence of Vps39 - Vps8 hybrid complex is possible. The formation of such complex could not be seen in MS data after anti-HA IP using fly-lysates expressing Vps8-9XHA from its endogenous promoter (Lőrincz et al. 2016).

We sought to investigate the possibility of hybrid complex formation in HEK293 cells. To this end, we expressed Vps39-FLAG and Vps8 or Vps39-FLAG and Vps41-HA (control for HOPS). After anti-FLAG IP we could detect Vps8 (Vps39 - Vps8 hybrid) or Vps41-HA (HOPS) in the eluates (Fig. 4A). Next, we used Tgfbrap1-FLAG together with Vps41-HA or with Vps8 (control for CORVET). In these experiments,we could detect only Vps8 in the anti-FLAG IP eluate (Fig. 4B). Based on these, we showed that human Vps39 - Vps8 hybrid complex can form, while we could not detect the Vps41 - Tgfbrap1 complex.

**Figure 4.**
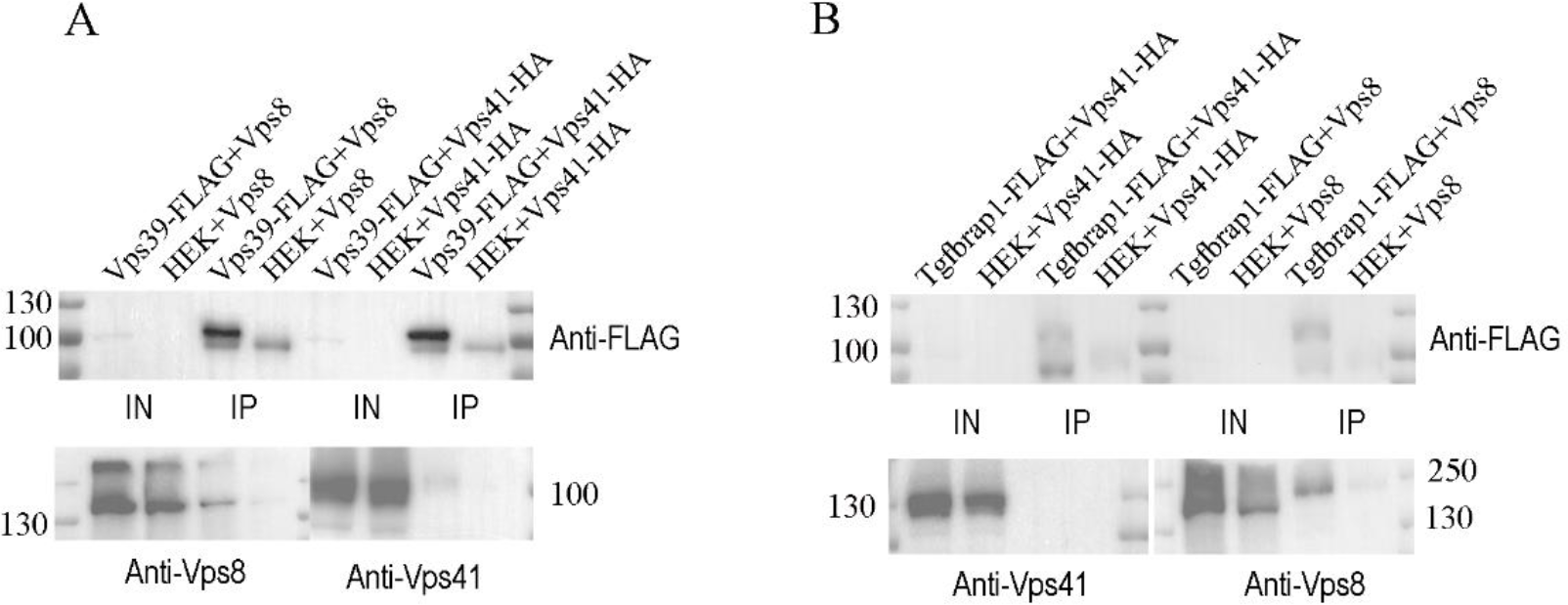
Hybrid complex containing Vps39 and Vps8 can form in HEK293 cells. Cells were transfected with the combination of Vps39-FLAG and Vps8 or with Vps39-FLAG and Vps41-HA (A). After anti-FLAG IP not just Vps41 (HOPS), but Vps8 (Vps39 – Vps8 hybrid) could be detected in the eluates. Transfection was also repeated with the following combinations: Tgfbrap1-FLAG and Vps41-HA or with Tgfbrap1-FLAG and Vps8 (B). Tgfbrap1 could bind only Vps8 (CORVET), Vps41 could not be detected in the eluates after anti-FLAG IP.

## Discussion

In our study, we investigated the assembly of two endosomal tethering complexes, namely CORVET and HOPS, which act in different stages of the endolysosomal system. We showed that overproduction of human CORVET-specific Vps8 or Tgfbrap1 decreased the amount of core subunits, Vps11 and Vps18 assembled with HOPS. In line with this, elevated amount of CORVET-specific subunits increased lipidated LC3 level and p62, in line with the expected failure in autophagosome-lysosome fusion due to defective HOPS function. The overexpression of HOPS-specific Vps41 or Vps39 did not affect either the amount of core subunits assembled with CORVET, or the level of LC3II or p62. We could detect that a hybrid complex, containing HOPS-specific Vps39 and CORVET-specific Vps8, can be formed in HEK293 cells.

HOPS and CORVET share four so-called Class C subunits (Vps33, Vps16, Vps8, Vps11) and their function and specificity are defined by Rab-binding complex-specific subunits. The existence of shared subunits suggests that cells must keep a fine balance between the levels of the two complexes and hence changes in the expression level of specific subunits may affect complex assembly. Of note, in yeast, CORVET-specific Vps3 overexpression caused vacuole fragmentation, which resembles to HOPS loss of function phenotype, and the level of HOPS also decreased (Peplowska et al. 2007). Additionally, yeast Vps39 and Vps3 occupy overlapping binding sites on Vps11 (Ostrowicz et al. 2010) and human Vps39 and Tgfbrap1 shares their binding sites on Vps11 as well (Van Der Kant et al. 2015). These altogether suggest competition between these specific subunits and are in line with our overexpression IP results, as elevated level of Tgfbrap1 decreased the amount of Vps11 and Vps18 that are bound with Vps39 (Fig.1B).

The overexpression of human Vps8, the other CORVET-specific subunit, has similar effect as Tgfbrap1, since the levels of Vps41-bound core subunits are decreased (Fig.1A). In yeast, overproduction of Vps8 did not affect HOPS function and the level of Vps41-bound (HOPS-assembled) Vps33 was unaffected as well (Peplowska et al. 2007). In contrast, overexpression of *Drosophila* Vps8, the only (mini)CORVET-specific subunit, inhibited all HOPS-dependent trafficking routes by outcompeting Vps41 from HOPS and decreased the level of Vps41-bound Vps16, Vps18 and Vps33 (Lőrincz et al. 2019). A smaller version of CORVET exists in flies, which lacks a Vps3 homolog (Lőrincz et al. 2016), that is why in this model only one of the CORVET-specific subunits can be examined. In human, both complexes are hexamers, and according to our results, both of the CORVET-specific subunits can compete with their HOPS-specific counterparts and as a consequence of their overproduction, less HOPS-specific core subunits could be detected (Fig.1A and B). This suggests that less HOPS can be assembled when any of the CORVET-specific subunits (Vps8 or Tgfbrap1) are overexpressed.

Next, we investigated if the surplus of HOPS-specific subunits, Vps39 and Vps41, affect the level of CORVET-bound core subunits. We found that despite the overproduced HOPS-specific subunits, the levels of the CORVET-bound (Tgfbrap1- or Vps8-assembled) Vps11 or Vps18 were comparable to control cells (Fig2A and B). Similar results were obtained in flies, since the overexpression of HOPS-specific Vps41 did not disturb CORVET-related functions (Lőrincz et al. 2019).

Our data suggest that CORVET-specific subunits may outcompete HOPS-specific ones, and hence they can affect HOPS function. Since HOPS mediates autophagosome-lysosome fusion (besides homotypic fusion of Rab7-positive structures), we determined the levels of lipidated LC3 and the autophagic cargo, p62. These proteins are accumulated on the autophagosomal membrane or in the autophagosome, respectively, if the fusion process is inhibited. We observed elevated LC3II levels in cells overexpressing Tgfbrap1 or Vps8 (CORVET-specific subunits), while we detected increased amount of p62 in Vps8 overproducing cells (Fig. 3A and B). These are in line with a failure of HOPS function. Similarly, accumulation of Ref(2)P/p62 and lipidated Atg8a is observed in *Drosophila* systematically overexpressing Vps8 (Lőrincz et al. 2019), while yeast cells with surplus of Vps3 showed fragmented vacuoles, resembling to cells lacking Vps39 (Peplowska et al. 2007). Taken together, overexpression of human Vps8 and Tgfbrap affect HOPS function, presumably by outcompeting their HOPS counterparts.

As both human complexes are hexamers and they have shared subunits, it allows the possible existence of such CORVET-HOPS hybrid complexes, just like in yeast cells. According to our data, human Vps39-FLAG could precipitate not just the HOPS-specific Vps41, but the CORVET-specific Vps8 as well, which indicates a possible formation of human hybrid complex containing Vps39 and Vps8 (Fig.4A). In contrast, Tgfbrap-FLAG could not pull down HOPS-specific Vps41, only the CORVET subunit Vps8 (Fig.4B). However, it should be mentioned that overexpressed proteins were used in these experiments. Earlier data show the existence of such hybrid complex in yeast as well, since Vam6/Vps39 can bring downVps8 (besides other HOPS subunits), however, the amount of Vps8 was much lower compared to its HOPS counterpart Vps41 (Peplowska et al. 2007). Additionally, in another study, only a pentamer complex could be identified from cells lacking Vps3 after TAP-purification via Vps8. This complex contained only Vps8 and core subunits: Vps11, Vps18, Vps33 and Vps16, but Vam6/Vps39 did not replace Vps3, even though they occupy the same binding sites on Vps11. In contrast, Vps41 replaces Vps8 in Vps8 mutant cells, and a hexamer Vps41-Vps3 hybrid complex forms (Ostrowicz et al. 2010). Hence, it seems that possible formation of both hybrid complexes are seen in yeast, but the stability and functionality of them are different –it is likely that the level of the Vps41-Vps3 hybrid complex is higher. The only possible hybrid complex (Vps8-Vps39) could not be identified in flies (Lőrincz et al. 2016), and in accordance with this, neither Vps39 could be observed in lysates of GTP-locked Rab5 expressing Drosophila S2 cells (CORVET binding partner), nor Vps8 in lysates of GTP-locked Rab2 expressing cells (HOPS binding partner) (Gillingham et al. 2014). Based on our results, in human cells the Vps39-Vps8 hybrid complex can be formed (Fig 4A).

## Materials and Methods

### Cloning

To obtain DNA constructs for stably transfected cell lines generation, our previous plasmid constructs were used as a template for amplification of the DNA regions encoding full-length proteins fused with FLAG- or HA-tags (Simon-Vecsei et al. 2021). The amplified DNA parts were cloned into mammalian pCMV3 expression vector (SinoBiologicals) to Acc65I - NotI restriction sites using NEBuilder HiFi DNA Assembly Cloning Kit. N-terminal tags were used in the case of Vps41 and Vps39, while C-terminal for Vps8 and Tgfbrap1. We used Vps8 without tag as well, where it is indicated. The sequences were checked with Sanger sequencing (Microsynth AG, Switzerland).

### Cell culture and transfection

HEK293 (human embryonic kidney) cells were maintained as earlier (Simon-Vecsei et al. 2021). Briefly, high glucose Dulbecco’s Modified Eagle Medium (DMEM, Lonza) supplemented with 10% (v/v) heat-inactivated foetal calf serum (FBS, Lonza), 2mM L-glutamine (Lonza), 100 U/ml penicillin and 100mg/ml streptomycin were used. Cells were grown in standard conditions (at 37°C in a humidified atmosphere with 5% CO2). For western blot and IP experiments, 1.5 million HEK293 cells were plated into T25 flasks and transfected after 24 hours. Transfection was performed using TransIT-LT-1 (Mirus) transfection reagent and 2500ng plasmid/flask, according to the manufacturer’s instructions.

For the generation of stably transfected cell lines, cells were selected with Hygromycin (in 200 μg/ml final concentration) for at least two weeks.

### Immunoprecipitation

IPs were performed according to our earlier studies (Simon-Vecsei et al. 2021), with the following modifications. 24 hours after transfection cells were scraped in cold phosphate buffer saline (PBS), then centrifuged and resuspended in lysis buffer (50mM TRIS-HCl, pH: 7,5, 150mM NaCl, 1% Triton X-100, 5mM EDTA, 1mM PMSF, protease inhibitor coctail (Roche)), then incubated on ice for 20 minutes. Samples were centrifuged and 20μl anti-FLAG beads (Sigma) were added to the lysates. Sample were rotated for 1.5 hours at 4°C. After centrifugation, beads were rotated with wash buffer (same as the lysis buffer without protease inhibitor) for 3 × 10 minutes at 4°C. Bound proteins were eluted with Laemmli buffer, boiled for 5 minutes at 100°C and analyzed by western blot.

### Western blot

Protein samples were made 24 hours after transfection. Cells were washed with ice-cold PBS, scraped on ice in lysis buffer (50 mM Tris-HCl pH 8.0, 50 mM KCl, 10 mM EDTA, 1 mM PMSF, 1% Triton X-100) and incubated for 16 minutes on ice. After centrifugation, the protein amounts of the supernatants were measured by Bradford reagent (Thermo Scientific). Samples were boiled with Laemmli buffer at 100°C for 5 minutes. 20μg protein per sample were applied on the SDS polyacrylamide gel and after electrophoresis proteins were transferred to PVDF membranes. The following primary antibodies were used: anti-FLAG (Sigma, M2, mouse, 1:2000), anti-HA (Roche, rat, 1:1000), anti-Vps18 (Abcam, rabbit, 1:3000), anti-Vps11 (Sigma, rabbit, 1:500), anti-Vps8 (Atlas, rabbit, 1:500), anti-Vps41 (Abcam, rabbit, 1:1000), anti-Tgfbrap1 (Abcam, rabbit, 1:200), anti-LC3 (Nanotools, mouse, 1:200), anti-p62 (Medical & Biological Laboratories, rabbit, 1:1000), anti-tubulin (DSHB, mouse, 1:800). For development of specific protein bands, horse-radish peroxidase (HRP) conjugated secondary antibodies were used in 1:2500 (anti-rabbit, DAKO) or in 1:10.000 (anti-mouse, Sigma) dilution. Protein detection were performed with Immobilon Western Chemiluminescent HRP Substrate (ECL, Millipore). Band intensities were analyzed by ImageJ software.

## Acknowledgements

We thank to Sarolta Pálfia for technical assistance.

This work was supported by the ÚNKP-22-3 New National Excellence Program of the Ministry for Culture and Innovation from the source of the National Research, Development and Innovation Fund. (ÚNKP-22-3-I-ELTE-547 to MM).

## Notes

### Competing Interest Statement

The authors have declared no competing interest.

